# Cooperative Effects of RIG-I-like Receptor Signaling and IRF1 on DNA Damage-Induced Cell Death

**DOI:** 10.1101/2021.10.21.465312

**Authors:** David Y. Zander, Sandy S. Burkart, Sandra Wüst, Vladimir G. Magalhães, Marco Binder

## Abstract

Properly responding to DNA damage is vital for eukaryotic cells, including the induction of DNA repair, growth arrest and, as a last resort to prevent neoplastic transformation, cell death. Besides being crucial for ensuring homeostasis, the same pathways and mechanisms are at the basis of chemoradiotherapy in cancer treatment, which involves therapeutic induction of DNA damage by chemical or physical (radiological) measures. Apart from typical DNA damage response mediators, the relevance of cell-intrinsic antiviral signaling pathways in response to DNA breaks has recently emerged. Originally known for combatting viruses via expression of antiviral factors including IFNs and establishing of an antiviral state, RIG-I-like receptors (RLRs) were found to be critical for adequate induction of cell death upon the introduction of DNA double-strand breaks. We here show that presence of IRF3 is crucial in this process, most likely through direct activation of pro-apoptotic factors rather than transcriptional induction of canonical downstream components, such as IFNs. Investigating genes reported to be involved in both DNA damage response and antiviral signaling, we demonstrate that IRF1 is an obligatory factor for DNA damage-induced cell death. Interestingly, its regulation does *not* require activation of RLR signaling, but rather sensing of DNA double strand breaks by ATM and ATR. Hence, even though independently regulated, both RLR signaling and IRF1 are essential for proper induction/execution of intrinsic apoptosis. Our results not only support more broadly developing IRF1 as a biomarker predictive for the effectiveness of chemoradiotherapy, but also suggest investigating a combined pharmacological stimulation of RLR and IRF1 signaling as a potential adjuvant regimen in tumor therapy.

## Introduction

DNA damage is a ubiquitous and existential threat to organisms. Potential causes comprise ionizing radiation (IR), genotoxic chemicals, but also cell-intrinsic mechanisms. Among various possible DNA alterations, the most drastic and impactful are DNA double-strand breaks (DSBs). Complex mechanisms involving detection by ATM, ATR, and downstream processes including the tumor suppressor p53 and checkpoint inhibition, either lead to sufficient repair of the damage or to induction of programmed cell death [1, 2]. The latter mostly comprises apoptosis, but other forms such as necroptosis and pyroptosis have recently been reported as well. Mutations of the central DSB sensors can cause severe diseases such as ataxia telangiectasia, associated with carcinogenesis and serious immunodeficiency [3–5]. Originally discovered and best-studied in the context of the antiviral innate immune response, IRF1 has been implicated in the DNA damage response and tumor suppressor functions [6–9].

Following the IRF1 example, it became apparent that cell-intrinsic antiviral signaling pathways also substantially contribute to DNA damage-induced cell death. Both STING and RIG-I-like receptor (RLR) pathways detect damage-associated molecular patterns (DAMPs), such as endogenous DNA fragments and nuclear RNA, and can trigger cell death [10, 11]. Previously, RIG-I stimulation has been shown to induce death of breast cancer cells, putting forward a potential application in tumor therapy [12]. Typically, the RLRs, RIG-I and MDA5, are stimulated by non-self RNA in the event of viral infection. Interaction with their adaptor MAVS leads to activation of the transcription factors IRF3, NF-κB p65/RELA and p50/NFKB1. The resulting expression of ISGs and IFNs of type I/III causes the establishment of an antiviral state and, in most cases, effective containment of the invading pathogen. In addition to apoptosis sensitizing effects of NF-κB and IFNs through expression of pro-apoptotic factors, direct cell death mediating effects have recently been reported for MAVS and IRF3 [13, 14]. Chattopadhyay et al. were first to identify and characterize the RLR-induced IRF3-mediated pathway of apoptosis (RIPA) [15]. Stimulation of RLRs with dsRNA or viral infection induces MAVS-dependent ubiquitination of IRF3 and subsequent activation of pro-apoptotic factors independent of IRF3’s transcriptional activity [16]. Furthermore, MAVS was shown to directly interact with procaspase-8, forming so-called MAVS-death-inducing signaling complexes upon viral infection [17]. Here we show that RLR signaling, IRF1, and canonical DNA damage response pathways, comprising ATM/ATR and p53, are essential for efficient induction of apoptosis. We show that these pathways have independent pro-apoptotic capacities, and we present new insights into IRF1’s complex cellular functions.

## Methods

### Cell culture, cell line generation, and stimulation

Cell lines were grown at 37 °C, 95 % humidity, and 5 % CO_2_ in Dulbecco’s modified eagle medium (DMEM high glucose, Life Technologies, Carlsbad, CA, USA), supplemented with final 10 % (v/v) fetal calf serum (FCS, Thermo Fisher Scientific, Waltham, MA, USA), 1x non-essential amino acids (Thermo Fisher Scientific), and 100 U/ml penicillin and 100 ng/ml streptomycin (LifeTechnologies). For generation of transgene expressing A549 cell lines by lentiviral transduction, lentiviral particles were produced by transfecting HEK 293T cells with plasmids pCMV-dr8.91, pMD2.G, and the respective retroviral vector (pWPI) using calcium phosphate transfection (CalPhos Mammalian Transfection Kit, Takara Bio Europe, Saint-Germain-en-Laye, France). After two days the supernatant was harvested, sterile filtered, and used to transduce target cells two times for 24 h. Transduced cells were selected with antibiotics appropriate for the encoded resistance gene (5 μg/ml blasticidin, MP Biomedicals, Santa Ana, CA, USA; 1 μg/ml puromycin, Sigma Aldrich; 1 mg/ml geneticin (G418), Santa Cruz, Dallas, TX, USA). Knockout (KO) cell lines were generated by clustered regularly interspaced short palindromic repeats (CRISPR)/Cas9 technology. DNA oligonucleotides coding for guideRNAs against the respective genes (sequences shown in Supplementary Table S1) were cloned into the expression vector LentiCRISPRv2 (Feng Zhang, Addgene #52961).

Transduced A549 wild-type cells were selected with puromycin, single cell clones were isolated, and KO was validated by immunoblotting and functional tests (Fig. S5). A549 *IFNAR1*^-/-^ *IFNLR1*^-/-^ *IFNGR*^-/-^ (IFNR TKO), *IRF1*^-/-^, *IRF1* OE, *IRF3*^-/-^, IRF3-eGFP H2B-mCherry, *MAVS*^-/-^, *MYD88*^-/-^*, RELA*^-/-^, and *RIG-I*^-/-^ were reported previously [18–22]. A549 *RIG-I* OE cells were generated by stable lentiviral transduction as described previously [19]. Cells transduced with non-targeting gRNA (sequence taken from the GeCKO CRISPR v2 library) were used as controls. PH5CH non-neoplastic hepatocytes and HepG2 cells were kindly provided by Dr. Volker Lohmann (Heidelberg University, Heidelberg, Germany). Huh7.5 cells were generously provided by Dr. Charles Rice (Rockefeller University, New York).

Stimulation was performed with doxorubicin (DOX, Hölzel Diagnostika, Cologne, Germany), etoposide (ETO, Cell Signaling Technology, Danvers, MA, USA), or cells were transfected with *in vitro* transcribed and chromatographically purified 200 bp 5’ppp-dsRNA [23], poly(C) (Sigma-Aldrich), and poly(I:C) (Sigma-Aldrich) using Lipofectamine 2000 (Invitrogen, Carlsbad, CA, USA) following the manufacturer’s protocol. Cells were γ-irradiated with doses of 0-30 Gy using a Gammacell 40 Exactor (Best Theratronics, Ottawa, Canada).

### Real-time imaging of cell death

A549 cells stably expressing histone H2B mCherry [21] were seeded at density of 2 x 10^3^ cells per 96-well. The next day, cells were stimulated with 1-2 μM DOX (10 h), 25 μM ETO (10 h), 0.1 ng/ml dsRNA (8 h), or γ-IR. DMSO (Carl Roth, Karlsruhe, Germany), poly(C), and mock irradiation were used as appropriate controls. Post treatment, fresh medium was supplemented with 1:10 000 IncuCyte^®^ Cytotox Green Reagent (Sartorius, Göttingen, Germany) to determine dead cells. Total cell number and dead cells were monitored every 2 h using a 10x magnification in an IncuCyte^®^ S3 Live-Cell Analysis System (Satorius, Göttingen, Germany). For IFN pre-stimulation, 200 IU/ml IFN-β (IFN-β1, Bioferon, Laupheim, Germany) or IFN-γ (R&D Systems, Minneapolis, MN, USA) were added at the time of seeding. For inhibitor administration, 40 μM Z-VAD-FMK (Z-VAD, R&D Systems) and 10 μM Necrostatin-7 (Nec-7, Sigma Aldrich), or 25 μM TPCA-1 (Sigma Aldrich) were added 2 h prior treatment. IncuCyte^®^ Software (2019B Rev2, Satorius, Göttingen, Germany) was used to mask cells in phase contrast images. Calculations were performed applying the following settings: red fluorescence: segmentation top-hat, radius 100 μM, threshold (GCU) 0.4, edge split sensitivity −35, area 60-1000 μm^2^, integrated intensity ≥ 60; green fluorescence: segmentation top-hat, radius 100 μM, threshold (GCU) 10, edge split sensitivity −40, area 100-700 μm^2^, eccentricity ≤ 0.8, mean intensity 7-1000, integrated intensity ≥ 2500. Percentage of dead cells was calculated relative to total cell count. Data represent the results of at least three biologically independent experiments. For curve charts, results were normalized to the control cell line of each replicate. Bars represent non-normalized means 36 h post treatment.

### Immunofluorescence microscopy and determination of cellular IRF3 distribution

Fluorescence microscopy was performed to visualize phosphorylated histone H2A.X. After 4 h treatment with 2 μM DOX or DMSO, or 1 h post γ-IR with 20 Gy or 0 Gy, cells were permeabilized with −20 °C methanol and fixed with 4 % paraformaldehyde. To block non-specific background, cells were incubated with 1 % (w/v) bovine serum albumin (BSA) and 10 % (v/v) FCS for 30 min. Primary antibodies specific for phospho-H2A.X (Cell Signaling Technology, 9718, 1:1000) were applied at 4 °C over-night. Slides were incubated with Alexa Fluor^®^ 488 anti-rabbit (ThermoFisher Scientific, Waltham, MA, USA, A11008, 1:1000) and DAPI (ThermoFisher Scientific, D1306, 1:5000) for 1 h. For determination of cellular IRF3 distribution, A549 cells stably expressing IRF3-eGFP and histone H2B-mCherry were stimulated either with DOX or poly(I:C) for 12 h. Fluorescence was visualized using a Primovert microscope (Carl Zeiss, Jena, Germany).

### Immunoblotting

Stimulated cells were lysed in Laemmli sample buffer, and digested with Benzonase^®^ Nuclease (Merck Millipore, Burlington, MA, USA). For inhibitor administration, 20 μM KU-55933 (Sigma-Aldrich), 25 μM Rabusertib (Hölzel Diagnostika), 25 μM TPCA-1 (Sigma Aldrich), or 10 μM VE-822 (Hölzel Diagnostika) were added 2 h prior treatment. For stimulation with IFNs, 200 IU/ml IFN-α (PBL Assay Science, Piscataway, NJ, USA), IFN-β, or IFN-γ were applied over-night. Lysed samples were further denatured at 95 °C for 5 min and cleared from detritus. Resulting protein extracts were subjected to 10 % (w/v) SDS-polyacrylamide gel electrophoresis and transferred to PVDF membranes (Bio-Rad, Hercules, CA, USA, 0.2 μm pore size). Upon incubation with 5 % (w/v) BSA for 2 h to block non-specific background, membranes were probed using antibodies specific for β-actin (Sigma-Aldrich, A5441, 1:5000), calnexin (Enzo Biochem, Farmingdale, NY, USA, ADI-SPA-865-F, 1:1000), CASP3 (Cell Signaling Technology, 9662S, 1:1000), CASP9 (Cell Signaling Technology, 9508, 1:1000), IRF1 (Cell Signaling Technology, 8478S, 1:1000), phospho-IRF3 (pS396, ThermoFisher Scientific, MA5-14947, 1:1000), JAK1 (Cell Signaling Technology, 3332S, 1:1000), MDA5 (Enzo Biochem, ALX-210-935, 1:1000), NFKB1 (p50) (Abcam, Cambridge, UK, ab32360, 1:1000), p53 (Santa Cruz Biotechnology, Dallas, TX, USA, sc-126, 1:1000), or STAT1 (BD Biosciences, Franklin Lakes, NJ, USA, 610115, 1:1000) at 4 °C over-night. For detection, anti-rabbit horseradish peroxidase (HRP) (Sigma-Aldrich, A6154-5X1ML, 1:20 000) or anti-mouse HRP (Sigma-Aldrich, A4416-5X1ML, 1:10 000) were applied for 1 h, membranes were covered with Amersham ECL Prime Western Blotting Detection Reagent (ThermoFisher Scientific) for 1 min, and luminescence was detected using a sensitive CCD camera system (ECL ChemoCam Imager 3.2, INTAS Science Imaging Instruments, Göttingen, Germany). Densitometric analysis of the protein bands was performed using ImageJ (1.52e). Data shown represent the results of at least three biologically independent experiments.

### Quantitative PCR with reverse transcription (qRT-PCR)

Upon stimulation, cells were lysed and total RNA was isolated with the Monarch RNA isolation kit (New England Biolabs, Ipswich, MA, USA), following the manufacturer’s protocol. After extraction, complementary DNA (cDNA) was generated using the High Capacity cDNA Reverse Transcription kit (ThermoFisher Scientific). Determination of messenger RNA (mRNA) expression was performed using iTaq Universal SYBR^®^ Green Supermix (Bio-Rad) on a CFX96 real-time-system (Bio-Rad). Sequences of specific exon-spanning PCR primers are shown in Supplementary Table S2. GAPDH mRNA was used as a housekeeping gene control and relative expression determined by 2^ΔCt^ (thus, not normalizing to reference condition).

### Cell Viability

A549 cells were seeded at a density of 6 x 10^3^ cells per 96-well. Upon treatment with 2 μM DOX or DMSO for 24 h, cell viability was determined using the CellTiter-Glo^®^ luminescent cell viability assay (Promega, Madison, WI, USA) following the manufacturer’s protocol. Luciferase activity was measured using a Mithras LB 943 multimode reader (Berthold Technologies, Bad Wildbad, Germany).

### Caspase activity

A549 cells were seeded at density of 6 x 10^3^ cells per 96-well. 48 h post treatment with 0-2 μM DOX for 10 h, caspase-3/7 activity was determined using the Apo-ONE^®^ homogeneous caspase-3/7 assay (Promega) following the manufacturer’s instructions. Resulting fluorescence was measured using the Mithras LB 943 multimode reader (Berthold Technologies).

### Statistics

Comparison of datasets was performed using a paired, two-tailed Student’s t-test. * indicates p ≤ 0.05, ** p ≤ 0.01, *** p ≤ 0.001, **** p ≤ 0.0001. Error bars represent standard deviation.

## Results

### Apoptosis induction via DNA damage response pathway in A549 cells

To investigate the molecular links between DNA damage-induced cell death and innate immune signaling, we used immunocompetent A549 human lung carcinoma cell lines with functional knockouts (KOs) of components of both pathways. Cells were treated with DNA DSB inducers, specifically γ-IR or the topoisomerase II inhibitors doxorubicin (DOX) and etoposide (ETO), and the resulting cell death was monitored on single-cell level by real-time imaging.

Treatment of A549 cells with DOX resulted in pronounced cell death (Fig. 1A) and a corresponding reduction of bulk cell viability (Fig. 1B), accompanied by the detection of the DNA damage marker phospho-histone H2A.X by immunofluorescence (Fig. 1C). As in DMSO control conditions no cell death was observed (Fig. 1A), for the clarity of presentation we omitted this control in the following figures (data was acquired in every experiment). In order to characterize the type of cell death predominant upon DOX-induced DNA damage, we first evaluated activation of caspase-3 and −7 being pivotal markers of apoptosis. DOX treatment activated caspase-3 and −7 in a dose-dependent manner (Fig. 1D). Conversely, we treated cells with the pan-caspase inhibitor Z-VAD, or depleted caspase-3 or −9. Both approaches resulted in a significant reduction of cell death upon DOX treatment (Fig. 1E, F, H). These findings confirmed prior reports that cell death driven by DOX is mainly due to apoptosis [24]. Next, we investigated typical components of the DNA damage response upstream of caspase activation. In line with p53’s (*TP53*) essential role in inducing apoptosis, depletion of p53 showed a significant reduction of cell death (Fig. 1G, H). Interestingly, *TP53*^-/-^ had the opposite effects at late time points, elevating cell death for time points >54 h (Fig. 1G). Amongst others, p53 induces apoptosis via activation of PUMA and NOXA. Accordingly, we found *PUMA* and *NOXA* transcript levels to be increased in DOX treated cells (Fig. 1I), supporting a canonical DNA damage response through p53 in DOX-treated A549 cells.

**Fig. 1.**
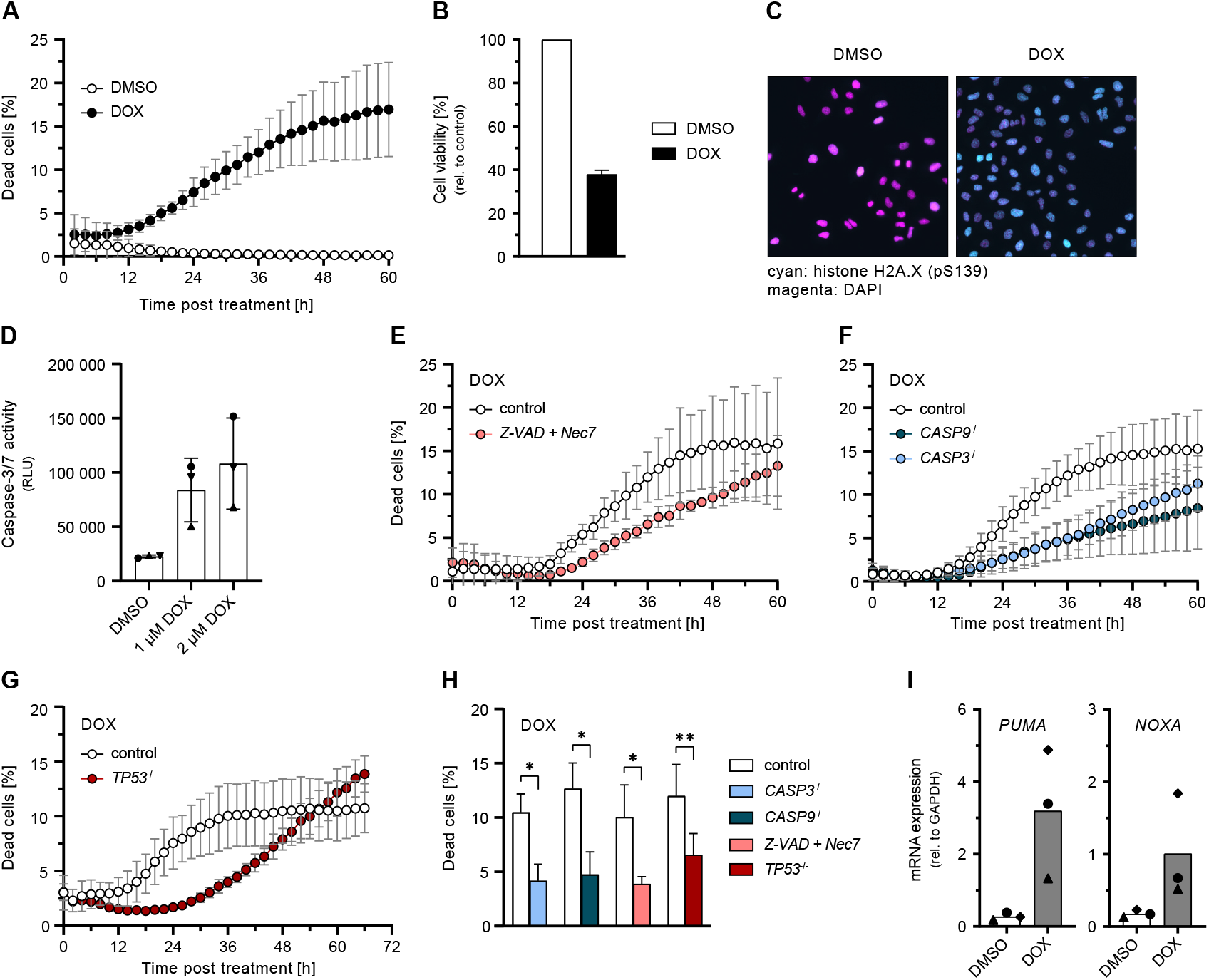
Induction of apoptosis upon DOX-mediated DNA damage. **(A**) Percentage of dead A549 cells relative to total cells counted over time post DOX or DMSO treatment. **(B)** Cell viability of A549 cells post DOX treatment for 24 h. **(C)** Immunofluorescence of phosphorylated histone H2A.X (S139) (cyan) and DAPI-stained nuclei (magenta) in A549 cells post DOX treatment for 4 h. **(D**) Caspase-3/7 activity of A549 cells 24 h post DOX treatment for 10 h. **(E-H)** Percentage of dead A549 cells with caspase inhibition or functional KO of the indicated genes relative to total cells counted over time **(E-G)** or 36 h **(H)** post DOX treatment. **(I)** A549 cells were treated with 1 μM DOX or DMSO for 24 h. PUMA and NOXA mRNA transcripts were determined by qRT-PCR. (**A, B, D-I**) Data shown represent the results of at least three biologically independent experiments.

### Relevance of innate antiviral immunity pathways in DNA damage induced cell death

In order to investigate the contribution of antiviral signaling cascades to the induction of DSB-induced cell death, we compared the impact of the major antiviral pathways using KOs of their respective signaling adapters. We observed DOX-induced cell death to be significantly reduced only by MAVS depletion (RLR signaling), but not so in the absence of STING (cGAS signaling), TRIF (TLR3 signaling), or MYD88 (general TLR signaling) (Fig. 2A–C). Despite RLR signaling appeared to play a major role, neither canonical IRF3 phosphorylation nor its nuclear translocation could be detected (Fig. 2D, E). Consistently, there was also no characteristic RLR-mediated induction of ISGs, such as *IFIT1* (Fig. 2F).

**Fig. 2.**
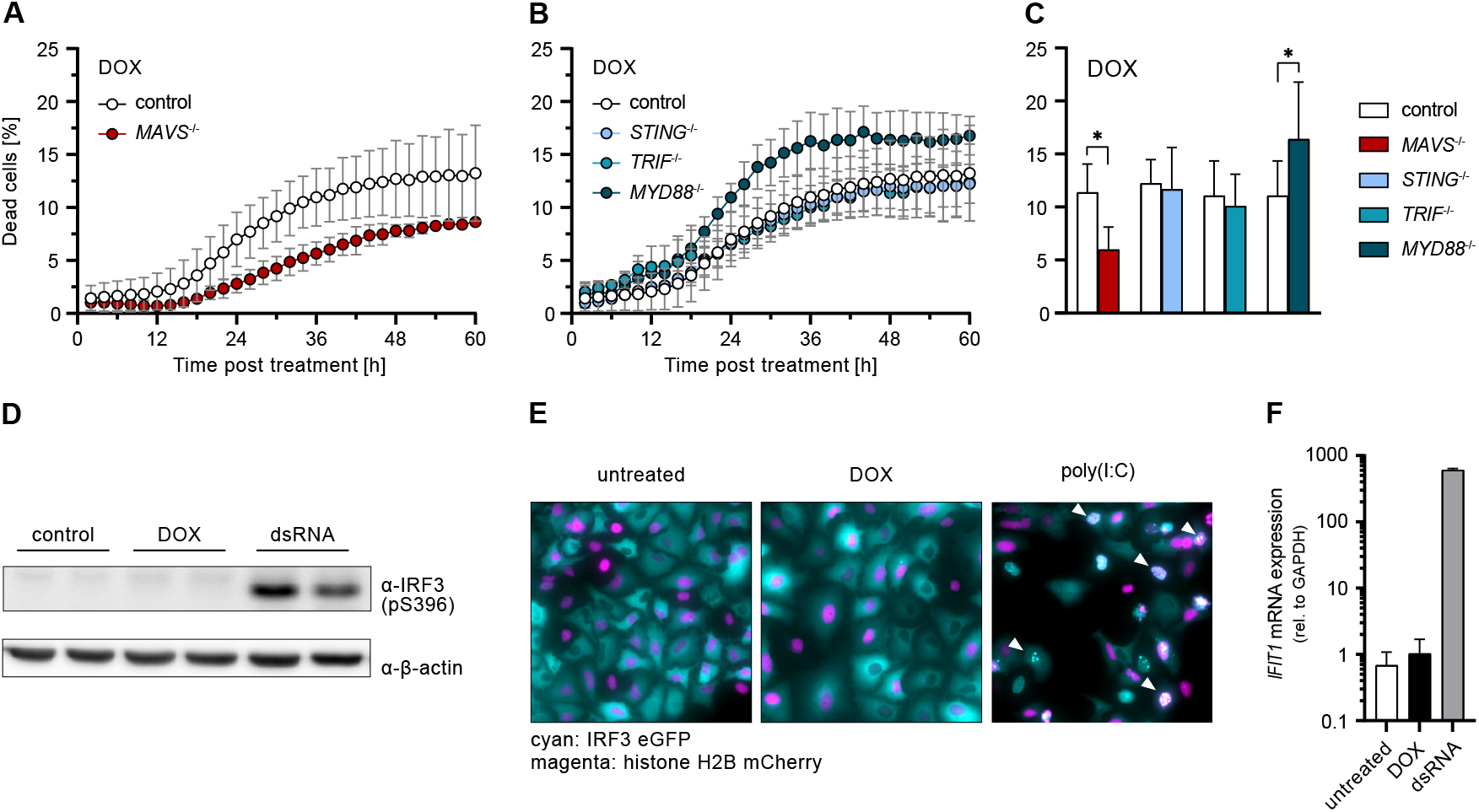
Relevance of antiviral signaling adapters and ISG response during DOX-induced DNA damage response. (**A-C)** Percentage of dead A549 cells with functional KO of the indicated genes relative to total cells counted over time **(A, B)** or 36 h **(C)** post DOX treatment. **(D)** A549 cells were stimulated with 1 μM DOX or 1 ng/ml dsRNA for 8 h. Phosphorylated IRF3 (S396) was determined by western blot. **(E)** A549 cells were stimulated with 1 μM DOX or 2 μg/ml poly(I:C) for 12 h. Cellular distribution of IRF3 eGFP (cyan) and histone H2B (magenta) was visualized by immunofluorescence microscopy. **(F)** A549 cells were stimulated with 1 μM DOX or 10 ng/ml dsRNA for 24 h. IFIT1 mRNA transcripts were determined by qRT-PCR. (**A-C, F**) Data shown represent the results of at least three biologically independent experiments.

Given the observed relevance of MAVS in DOX-induced cell death, we further analysed the effect of specific RLR depletion. Both *RIG-I*^-/-^ and *MDA5*^-/-^ reduced cell death upon DOX treatment, however, RIG-I exhibited a considerably stronger effect (Fig. 3A, C). Reciprocally, *RIG-I* overexpression (OE) markedly increased cell death upon DOX treatment (but not in untreated conditions, compare Fig. S1A), underlining the decisive role of RLR signaling in this process (Fig. 3B, C). In order to determine the factors responsible for mediating cell death downstream of MAVS, we further examined the influence of transcription factors IRF3 and NF-κB p65/RELA. We observed that depletion of either factor significantly reduced DOX-induced cell death (Figure 3D, F). Using IFN-“blind” A549 *IFNAR1*^-/-^ *IFNLR1*^-/-^ *IFNGR*^-/-^ (IFNR TKO) cells, we demonstrated that this effect was independent of a response mediated by secreted IFNs (Fig. 3E, F), which was further confirmed using *STAT1*^-/-^ cells (Fig. S1B). This was in accordance with the lack of ISG expression observed previously (Fig. 2F). Thus, IRF3 appears to have death sensitizing effects distinct from its classical transcriptional activity in the antiviral program.

**Fig. 3.**
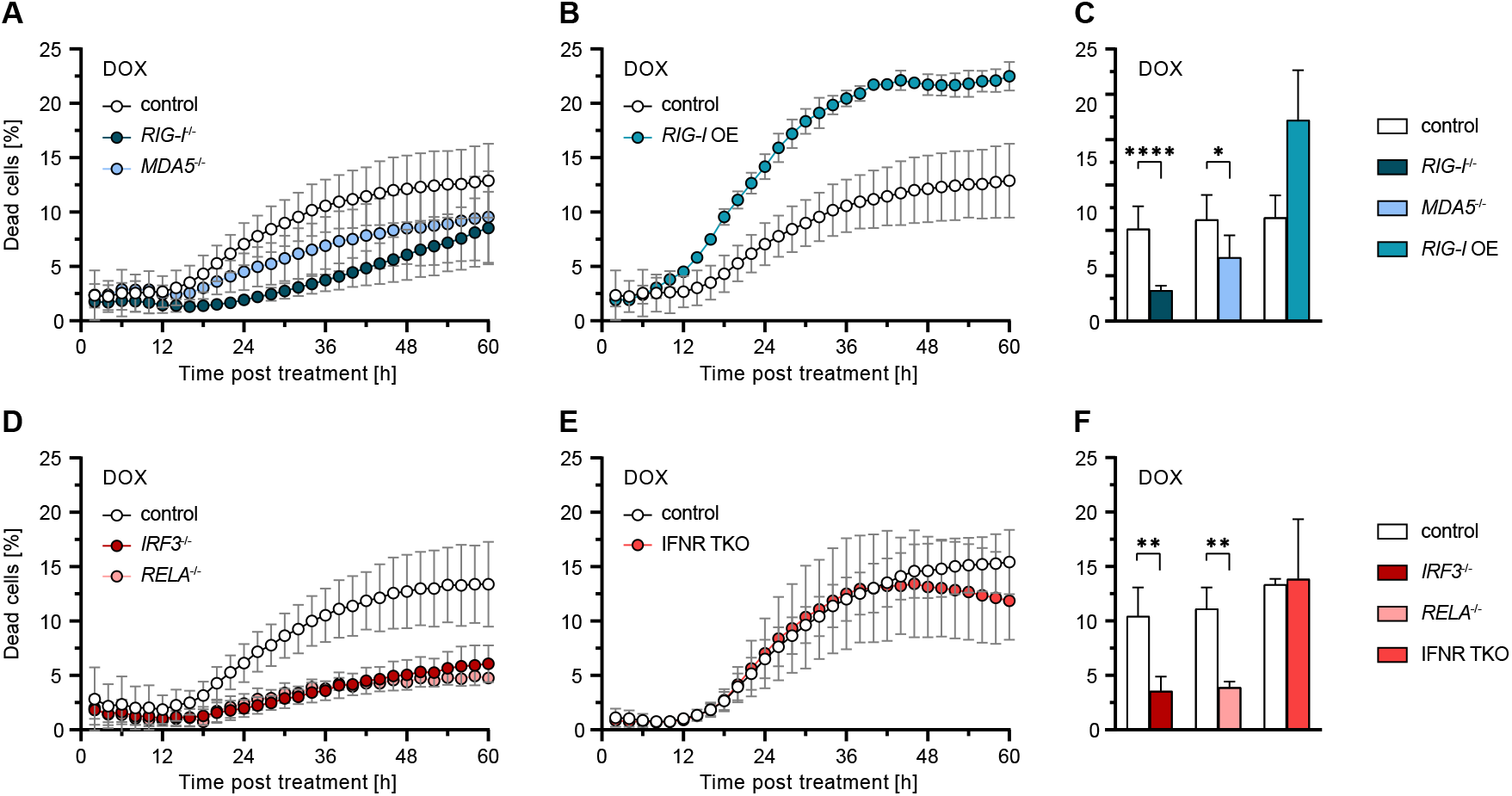
Implications of RLR signaling components and IFN signaling on DOX-induced apoptosis. **(A-F)** Percentage of dead A549 cells with functional KO or OE of the indicated genes relative to total cells counted over time **(A, B, D, E)** or 36 h **(C, F)** post DOX treatment. Data shown represent the results of at least three biologically independent experiments.

Taken together, we demonstrated that RLR signaling is required for the induction of cell death after DNA damage and that this function is independent of IFN secretion and the induction of canonical ISGs.

### Role of IRF1 in DNA damage induced apoptosis

Another transcription factor of the IRF family important for antiviral defenses [6, 18], IRF1, has previously also been implicated with the DNA damage response [25]. We hypothesized that upon genotoxic insult, IRF1 might be a downstream target of the RLR/IRF3 pathway, as reported for virus infection, and thereby link RLR activity to the DNA damage response. Indeed, upon DOX treatment, we observed IRF1 upregulation at the mRNA (Fig. 4A) and protein level (Fig. 4B). Of note, *IRF1* induction occurred independently of the presence of p53 (Fig. 4B). In order to determine the relevance of IRF1 to cell death, we next tested *IRF1^-/-^* cells in DOX treatment. Strikingly, IRF1 depletion almost completely abolished DOX-induced cell death (Fig. 4E, H). Conversely, increasing IRF1 abundance, either by OE through stable transduction or by pre-stimulation of cells with IFN-β or IFN-γ, markedly increased cell death upon DOX treatment (Fig. 4E, F, H), and the percentage of dead cells correlated with IRF1 levels in western blot (Fig. 4C, D). Notably, neither IFN stimulation alone, nor DOX treatment in IFN-primed but IRF1-depleted cells did induce cell death (Fig. S2A, B). Surprisingly, the same phenotype was observed in *RIG-I*^-/-^ conditions (Fig. S2C), in which IRF1 was present, suggesting a strict requirement of both RLR signaling and *IRF1* induction for proper triggering and/or execution of apoptosis. Similar observations were also made after ETO treatment (Fig. S2D, E), ruling out DOX-specific effects.

**Fig. 4.**
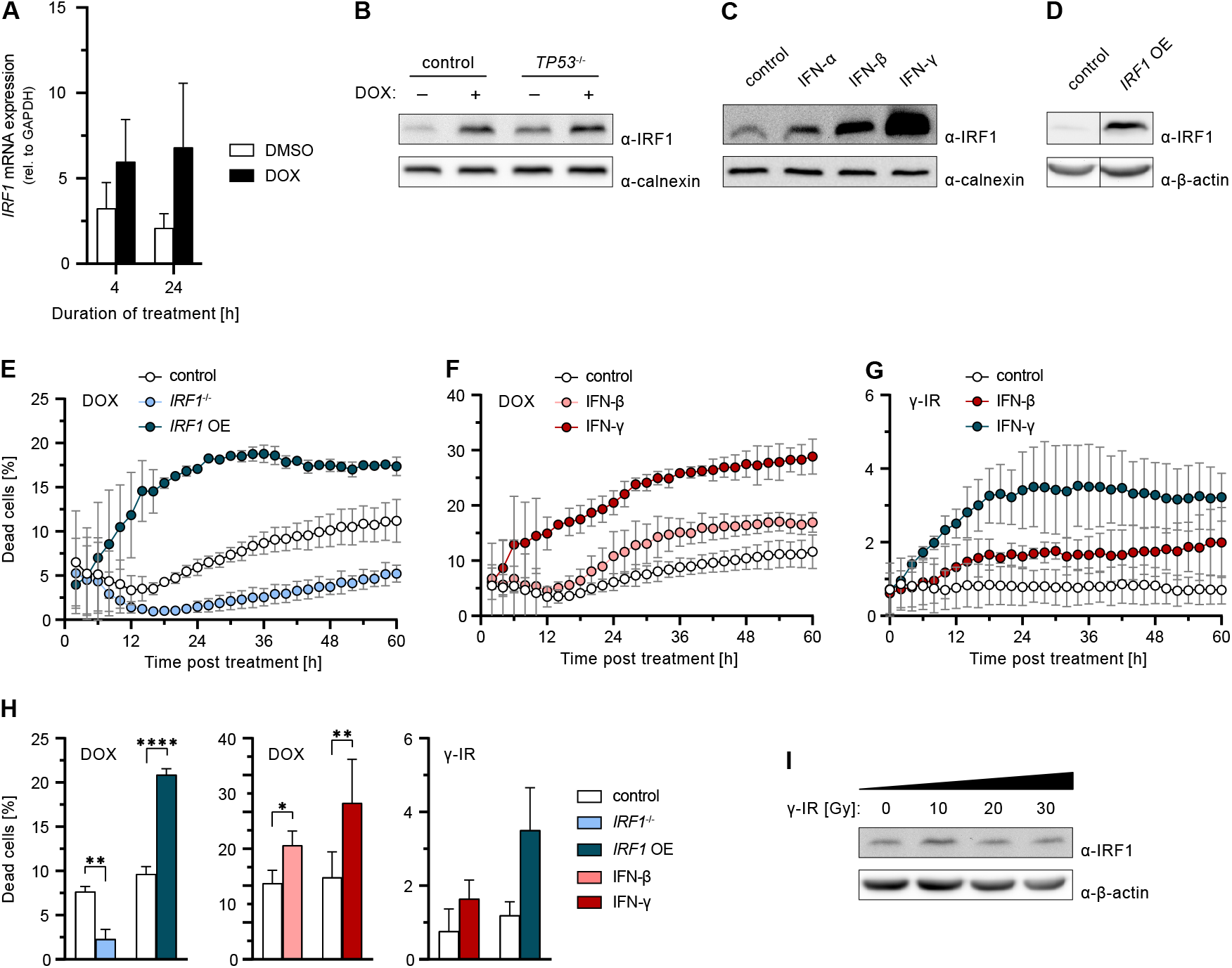
Relevance of IRF1 on DNA damage-induced cell death. **(A)** A549 cells were treated with 1 μM DOX or DMSO for 10 h. IRF1 mRNA transcripts were determined by qRT-PCR. **(B)** A549 cells were treated with 1 μM DOX or DMSO for 10 h. Levels of IRF1 were determined by western blot. **(C)** A549 cells were stimulated with IFN-α, IFN-β, or IFN-γ over-night. Levels of IRF1 were determined by western blot. (**D**) Levels of IRF1 in A549 control and *IRF1* OE cells were determined by western blot. **(E-H)** Percentage of dead A549 cells with functional KO or OE of IRF1, or post IFN pre-stimulation relative to total cells counted over time (**E-G**) or 36 h (**H**) post DOX or γ-IR (20 Gy) treatment. **(I)** A549 cells were γ-irradiated. After 10 h IRF1 protein levels were determined by western blot. (**A, E-H**) Data shown represent the results of at least three biologically independent experiments.

The fundamental importance of IRF1 was additionally demonstrated in response to γ-IR. Although irradiation did induce DNA damage in A549 cells (Fig. S2F), we could neither observe induction of *IRF1* expression nor any cell death upon administration of up to 30 Gy (Fig. 4G–I). Strikingly, induction of cell death upon γ-IR was restored under conditions of elevated IRF1 levels, such as stable OE or IFN-γ pre-stimulation (Fig. 4G, H). In line with this, cells in which γ-IR naturally leads to an upregulation of *IRF1* expression, such as PH5CH cells, did exhibit a dose-dependent induction of cell death (Fig. S2G, H).

Thus, we showed that besides p53 and RLR signaling, IRF1 is essential for proper triggering of cell death upon DNA damage. IFNs, in particular IFN-γ, sensitize cells for DNA damage-induced apoptosis through upregulation of IRF1.

### Regulation of *IRF1* expression upon DNA damage

Above we have shown that RLR/IRF3 signaling as well as expression of *IRF1* are crucially important for DNA damage-induced cell death. We further found IRF1 to be consistently induced under all tested conditions of DNA damage leading to cell death. We now aimed to confirm whether IRF1 is in fact induced as a downstream target of RLR signaling. We first investigated the induction of *IRF1* expression after RIG-I stimulation using dsRNA as a canonical, highly specific agonist [23]. Indeed, we observed a fully RLR-dependent (RIG-I, MAVS, IRF3) increase of IRF1 levels, with a partial contribution of p65/RELA and IFN signaling (IFNR TKO) (Fig. 5A), in line with a recent report of our lab [18]. dsRNA-stimulation furthermore also led to the induction of cell death, which was fully abolished upon depletion of the RLR signaling components RIG-I, MAVS, or IRF3 (Fig. 5B, D). Depletion of p65/RELA and the IFN receptors (IFNR TKO) had minor pro-survival effects, suggesting a major role for transcription-independent RIPA with a possible but limited role for IFN signaling and ISG induction (Fig. 5C, D). Interestingly and in clear contrast to the situation upon DNA damage, dsRNA-induced cell death was independent of IRF1 (Fig. 5C). Nonetheless, experimentally elevating IRF1 levels markedly increased the percentage of dead cells also in this setting (Fig. S3A, B).

**Fig. 5.**
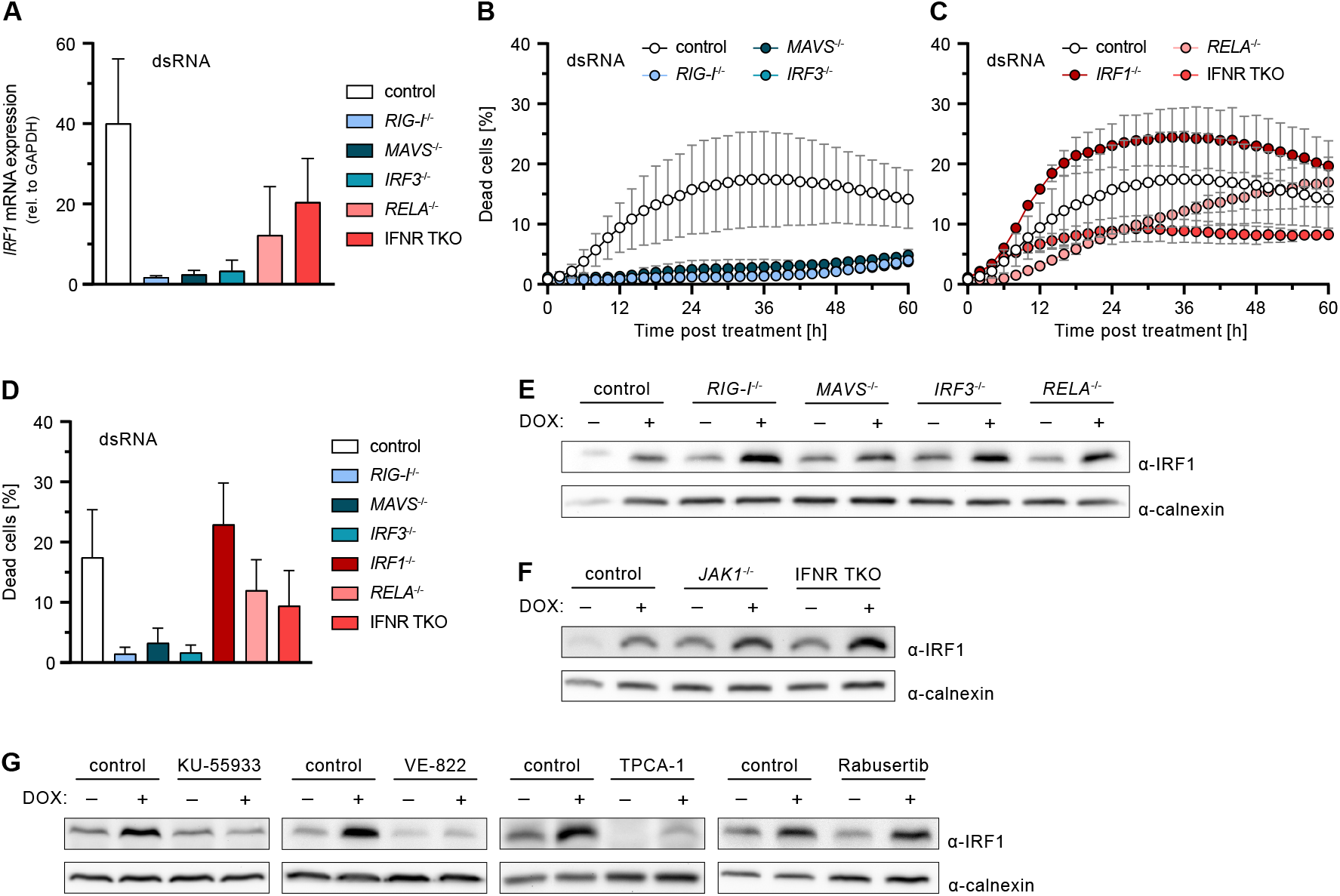
Effect of cell-intrinsic antiviral signaling components on dsRNA-induced cell death and *IRF1* expression. **(A**) A549 cells with functional KO of the indicated genes were stimulated with 2 ng/ml dsRNA for 6 h. IRF1 mRNA transcripts were determined by qRT-PCR. **(B-D)** Percentage of dead A549 cells with functional KO of the indicated genes relative to total cells counted over time **(B, C)** or 36 h **(D)** post dsRNA stimulation. **(E-G)** A549 cells with functional KO of the indicated genes or administration of the indicated inhibitors were treated with 2 μM DOX or DMSO for 6 h. Levels of IRF1 were determined by western blot. (**A-D**) Data shown represent the results of at least three biologically independent experiments.

These findings confirmed that, despite not being essential for cell death induction, IRF1 is induced downstream of RLR signaling, at least when stimulated by a strong RIG-I specific agonist. We next investigated whether this would be also the case in the context of DNA damage. Unexpectedly, upon treatment of cells with DOX, induction of *IRF1* expression was neither affected by depletion of RLR nor of IFN signaling components, including JAK1 (Fig. 5E, F; Fig. S3C). This suggested *IRF1* expression is induced independently of and coincidentally with antiviral RLR signaling upon DNA damage. We therefore hypothesized sensing of DNA damage might directly induce *IRF1*. To test this, we treated cells with specific inhibitors of the prototypical DSB sensors ATM and ATR, as well as potential downstream pathways. We found *IRF1* induction upon DOX-treatment to be completely blocked by the ATM inhibitor KU-55933 [26] and the ATR inhibitor VE-822 [27], suggesting important roles of these sensors in activation of IRF1 (Fig. 5G; Fig. S3D).

As *IRF1* expression has previously been shown to be NF-κB sensitive [28], we employed the common pan-NF-κB and JAK1 inhibitor TPCA-1 [29, 30]. Remarkably, TPCA-1 treatment completely prevented the induction of *IRF1* expression upon DOX treatment, and even strongly diminished basal expression (Fig. 5G, Fig. S3D). This effect could further be confirmed upon RLR-stimulation with dsRNA (Fig. S4A) and even upon IFN-γ treatment, which is a strong and well-studied canonical inducer of *IRF1* (Fig. S4B). We could rule out a cell line (A549) specific effect by testing three other human cell lines, PH5CH, HeLa and Huh7.5 (Fig. S4C). To our knowledge, this striking effect of TPCA-1 on *IRF1* expression has not been reported before. Again, corroborating IRF1’s crucial role in DNA damage-induced apoptosis, supressing *IRF1* induction by TPCA-1 also reduced cell death in DOX-treated A549, PH5CH, HeLa, and Huh7.5 cells (Fig. S4D).

Finally, we aimed to identify which signaling pathway and NF-κB subunit would be responsible for *IRF1* expression upon triggering the DNA damage response. As reported in literature, ATR may signal through CHK1 to activate p50/NFKB1, a potential target of TPCA-1 [31, 32]. We therefore inhibited CHK1 by Rabusertib [33] prior to DOX-treatment. However, our experiments did not reveal any effect of CHK1 inhibition or p50/NFKB1 depletion on IRF1 levels (Fig. 5G; Fig. S3D, E). We hence conclude that a so far elusive pathway downstream of the ATM/ATR system induces *IRF1*.

Taken together, we demonstrated that *IRF1* expression upon DOX-treatment is induced by the DSB sensors ATM/ATR rather than RLR signaling. This induction is independent of CHK1 signaling. Additionally, we identified a previously unappreciated IRF1-depleting effect of the NF-κB inhibitor TPCA-1.

## Discussion

Cells, particularly of multicellular organisms, have elaborate systems in place ensuring the integrity of their genome, as DNA damage poses severe risks of accumulating tumorigenic mutations or alterations. In response to excessive DNA damage beyond the potential of being properly repaired, cells trigger the execution of cell death programs, most commonly apoptosis [34]. This is also exploited for common cancer chemoradiotherapies, in which excessive DNA damage is radiologically (e.g., γ-IR) or pharmacologically (e.g., DOX or ETO) introduced, leading to the induction of cell death programs particularly in dividing tissues such as tumors. Elucidating the underlying mechanisms of how DNA damage molecularly leads to cell death is crucial to a better understanding of the circumstances leading to cancer and the pathways relevant for chemoradiotherapy. While classical DNA damage checkpoint control via p53 has been investigated thoroughly [1], much less is known about the relevance and contribution of non-canonical pathways. For example, a ground-breaking study surprisingly found the antiviral type I IFN pathway essential for certain chemotherapies’ efficacy [35]. Cytostatic and pro-apoptotic effects of IFNs have long been noticed [36–38]; however, it remained unresolved what triggered the production of IFNs in the studied context in the first place. Recent data also revealed cell-intrinsic triggering of cell death upon activation of antiviral signaling adapters, such as MAVS and STING. Interestingly, this was not only the case for viral infections, but also in response to DNA damage [10, 11, 39].

In the present study, we confirm this interrelationship between DNA damage response and antiviral signaling pathways, and we demonstrate an almost complete dependence of DOX- and ETO-triggered cell death on the presence of intact RLR/MAVS signaling. In clear contrast to recently published data, other branches of the cell-intrinsic antiviral defense, such as the TLR or the cGAS/STING system [10, 40, 41], did not affect DOX-induced cell death in our experimental setup. Instead, the cytosolic RNA sensors RIG-I and, to a lesser extent, MDA5 were triggered and essential for the induction of cell death. This is in line with a study by Ranoa et al. suggesting small nuclear RNAs U1 and U2 translocate into the cytoplasm in irradiated cells and trigger RIG-I activation [11]. In our experimental system, an intact RIG-I/MDA5-MAVS-IRF3 axis was essential for DNA damage induced cell death; however, we could not observe canonical transcriptional activities of IRF3, such as the induction of IFN genes or ISGs. While the relevance of both IRF3 and p65/RELA suggested the involvement of *IFNB* expression, KO of the receptors for all three types of IFNs (IFNR TKO) did not impact cell death. A plausible mechanism for this IFN-independent triggering of apoptosis is RIPA, involving LUBAC-dependent ubiquitylation of IRF3 and subsequent activation of pro-apoptotic BH3-only proteins [16]. The clear contribution of p65/RELA in our experiments might be through its transcriptional activation of further pro-apoptotic proteins [42]. To our knowledge, cooperative effects between RIPA and NF-κB have not been described before and may be an interesting subject for future investigations.

Efficient sensing of nuclear DSBs and triggering an appropriate response is critical for cell survival upon DNA damage, or for initiating cell death and preventing potentially cancerous transformation. As expected, we observed an essential role for p53, highlighting its central function in checkpoint control, coordinating DNA damage repair and triggering apoptosis as a last resort [43]. Interestingly, depletion of p53 reduced the number of apoptotic cells at early time points, but increased cell death at later times. Thus, absence of p53 led to a lack of induction of apoptosis in response to DOX-mediated DSBs at first, but likely massive accumulation of unrepaired DNA damage eventually led to increased, putatively necrotic cell death [44]. As a factor potentially linking the DNA damage response and antiviral signaling, we investigated the role of the multifunctional transcription factor IRF1, as it is known to be involved in both the DNA damage response [8, 25] and IFN signaling [6, 18, 45]. Indeed, we found that *IRF1* was considerably upregulated upon DOX and ETO treatment as well as γ-IR in different cell lines. Interestingly, only in A549 cells, described to be relatively radioresistant as a common characteristic for non-small cellular lung cancers [46], *IRF1* was not appreciably induced upon irradiation. We also observed a reduced histone H2A.X phosphorylation after γ-IR compared to DOX treatment, but potential underlying mechanisms are only partially understood and may comprise several processes [47, 48]. Nonetheless, we could further corroborate this clear correlation between *IRF1* induction and triggering/execution of a cell death program on a functional level. Experimentally increasing IRF1 levels by stable OE or by pre-treatment of cells with IFN-γ, known as a strong inducer of *IRF1* [45], radioresistance of A549 cells could be overcome. A similar effect has previously been demonstrated in T cells [25]. In our experiments, increased *IRF1* expression also led to a sensitization towards DOX-treatment. *Vice versa, IRF1* KO almost completely rescued cell survival upon DOX-, ETO- and γ-IR-induced DNA damage. These observations clearly establish a fundamentally important role of IRF1 in DNA damage-induced cell death. This is in accordance with literature suggesting IRF1 as a biomarker for radioresistance in tumor cells [49]. For example, extremely radioresistant osteosarcomas were shown to exhibit significantly reduced *IRF1* expression levels [50]. Our data further support establishing IRF1 as a predictive biomarker in chemoradiotherapy in tumor patients.

Our finding strongly suggested IRF1 to be the functional link between the DNA damage response and the antiviral system, with RLR signaling (either directly or via the IFN/JAK/STAT cascade) leading to transcriptional activation of IRF1. However, KO experiments clearly refuted this hypothesis. Neither KO of essential factors of the RLR pathway nor of IFN signaling components abolished *IRF1* induction upon DNA damage, suggesting that RLR signaling may activate IRF1 post-translationally. Generally, IRF1 is thought to be only regulated on a transcriptional level [45]. However, one study reports the requirement for “licensing” of IRF1 to become fully active, which required TLR signaling and MYD88 [51]. In preliminary experiments, we did not find any evidence for post-translational modifications in our setting, but this may warrant deeper investigations in the future. Alternatively, IRF1 might enhance the transcriptional response of IRF3, as reported before [52]. While we cannot rule out this possibility, the virtually complete inhibition of cell death in *IRF1*^-/-^ despite abundant presence of IRF3 makes this unlikely. In another study, we have also not found any indication of a dampening of IRF3 responses in A549 *IRF1*^-/-^ cells [18]. Notably, despite IRF1 being critically important for cell death induction in our system, *IRF1* (over-)expression alone did not suffice to elicit apoptosis. We therefore suspect RLR signaling and IRF1 activity to cooperate further downstream, putatively via the transcriptional activation of complementary pro-apoptotic factors.

It is interesting to note that cell death is also elicited upon RLR stimulation by dsRNA (the canonical way to trigger antiviral signaling). Also in this case, *IRF1* is induced, but strictly dependent on RIG-I and to a lesser extent dependent on IFN signaling. Surprisingly, however, depletion of IRF1 did not affect the cell death rate upon dsRNA stimulation, pointing towards transcription-independent mechanisms such as RIPA [15]. Still, KO of NF-κB (*RELA*) or the IFN receptors (IFNR TKO) affect cell death, suggesting some transcriptional regulation, which, however, was independent of IRF1. This may suggest that full-fledged RLR signaling upon dsRNA encounter induces a sufficiently broad transcriptional response, which (in contrast to the situation upon DNA damage) itself is capable of triggering apoptosis. Strikingly, even in dsRNA stimulation, ectopic OE of *IRF1* or pre-treatment of cells with IFN-γ led to a notable increase in the number of dying cells, putatively by the same cooperative pro-apoptotic effects observed in the case of DNA damage. This observation of a general sensitization for cell death by IRF1 is in line with data showing that *IRF1* OE enhances apoptosis in breast or gastric cancer treatment [53–55]. It is further plausible to speculate that reported pro-apoptotic effects of type I IFN [56, 57] would also be mediated by upregulation of *IRF1* through homodimeric STAT1 transcription factor complexes (GAF) inadvertently formed early upon IFNAR engagement [58]. This could mechanistically explain how IFN-α improved chemotherapy response and overall survival in a murine tumor model [35]. Thus, evidence further accumulates suggesting *IRF1*-inducing agents to be more broadly considered as adjuvants in tumor therapy.

Two central questions remain: firstly, which pro-apoptotic factors are specifically induced by IRF1 upon DNA damage that so potently sensitize cells to committing suicide upon (slight) RLR triggering. To this end, we are currently investigating IRF1-dependent candidate genes induced upon DOX-treatment at a transcriptomic level. Secondly, how is *IRF1* induced upon DNA damage in the first place if not through classical STAT1:STAT1 activity. In our study, we found its transcriptional regulation to be fully independent of RLR signaling and p53 but completely reliant on DNA DSB sensing via ATM and ATR. Still, the downstream pathway leading to *IRF1* expression remains elusive. While p65/RELA or p50/NFKB1 depletion did not affect *IRF1* induction, it was completely abolished by TPCA-1, a commonly known inhibitor of NF-κB. Interestingly, TPCA-1 considerably reduced baseline *IRF1* expression independent of the cell line used, and could even abolish the strong induction upon IFN-γ treatment. Thus, in addition to its inhibitory effects on NF-κB, JAK1, and STAT3 [29, 30, 59], TPCA-1 appears to specifically and very efficiently inhibit the activity of an essential transcription factor for *IRF1*.

In conclusion, our study highlights the critical relevance of the antiviral RLR system for the proper and timely induction of cell death upon DNA damage. We provide evidence for independent but cooperative involvement of p53, IRF1 and IRF3 activity upon detection of DNA DSBs by the ATM/ATR machinery. We show that elevating expression levels of *IRF1* lead to the sensitization towards cell death across different genotoxic insults, such as chemotherapeutics, γ-IR or cytosolic dsRNA (i.e. virus infection). These data corroborate a fundamental role for IRF1 and RLR signaling in DNA damage-mediated cell death and suggest future exploration of *IRF1* inducers, such as IFN-γ, together with low-dose RIG-I agonists for their potential as highly efficacious adjuvants in chemoradiotherapy. Additionally, our findings support IRF1 as a biomarker predictive for chemo- and radio-sensitivity of tumors.

## Supporting information

Supplementary Figures (with legends)

## Acknowledgements

We want to thank Joschka Willemsen and Leanne Strauß for having started our lab’s research line on RLR-related cell death and providing valuable preliminary data, Hendrik Welsch for help with the IncuCyte instrument and fruitful discussions, Volker Lohmann and Charlie Rice for provision of cell lines, and Ralf Bartenschlager for providing an excellent research environment and supervising the doctoral thesis of D.Z.. We are grateful to Eva Schnober, Hartmut Hengel and the rest of the team for managing the fantastic Integrated Research Training Group “Immunovirology” as part of TRR179, supporting D.Z. with a doctoral fellowship.

## Author contributions

This study was conceived and designed by M.B. and D.Z., critical intellectual input was provided by V.G.M., experiments were performed by D.Z. with assistance and contributions by S.W., S.S.B. and M.G.V., data was evaluated by D.Z. and M.B., the manuscript was written by D.Z. and M.B. and edited and approved of by all authors.

## Ethics Approval

This research did not involve human or animal material; ethical approval was not required.

## Data Availability

The raw data acquired for this study are available from the corresponding author on reasonable request.

