## Supplementary Figures (with legends) for "Cooperative Effects of RIG-I-like Receptor Signaling and IRF1 on DNA Damage-Induced Cell Death"

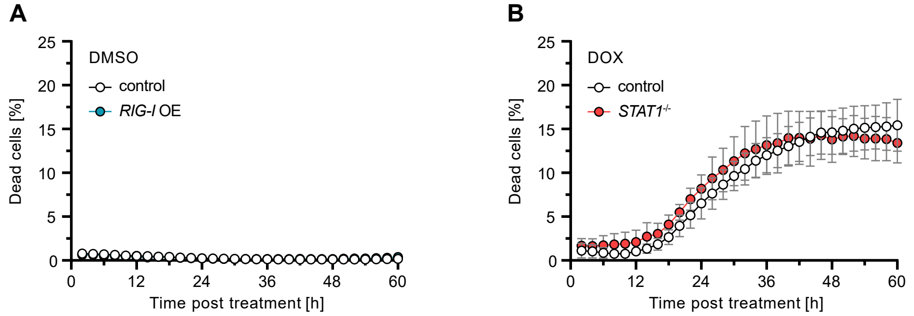


**Fig. S1.**

**(A, B**) Percentage of dead A549 cells with functional KO or OE of the indicated genes relative to total cells counted over time post DMSO or DOX treatment.

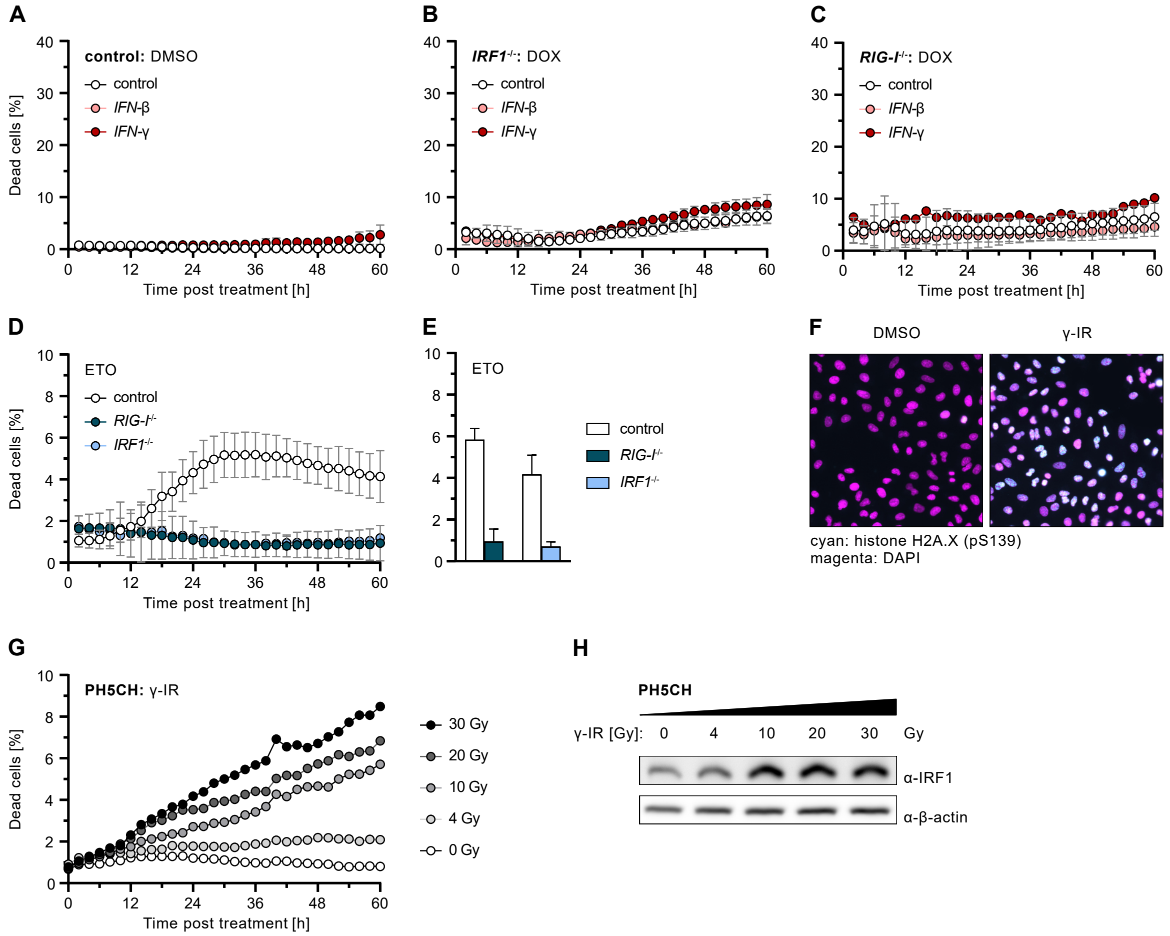


**Fig. S2.**

**(A-C)** Percentage of dead A549 cells with functional KO of the indicated genes relative to total cells counted over time post IFN pre-stimulation and DOX treatment. **(D, E)** Percentage of dead A549 cells with functional KO of the indicated genes relative to total cells counted over time **(D)** or 36 h **(E)** post ETO treatment. **(F)** Immunofluorescence of phosphorylated histone H2A.X (S139) (cyan) and DAPI-stained nuclei (magenta) in A549 cells 1 h post γ-IR. **(G)** Percentage of dead PH5CH cells relative to total cells counted over time post γ-IR. **(H)** PH5CH cells were treated with γ-IR at different doses. After 8 h IRF1 protein levels were determined by western blot. (**A-E**) Data shown represent the results of at least three biologically independent experiments.

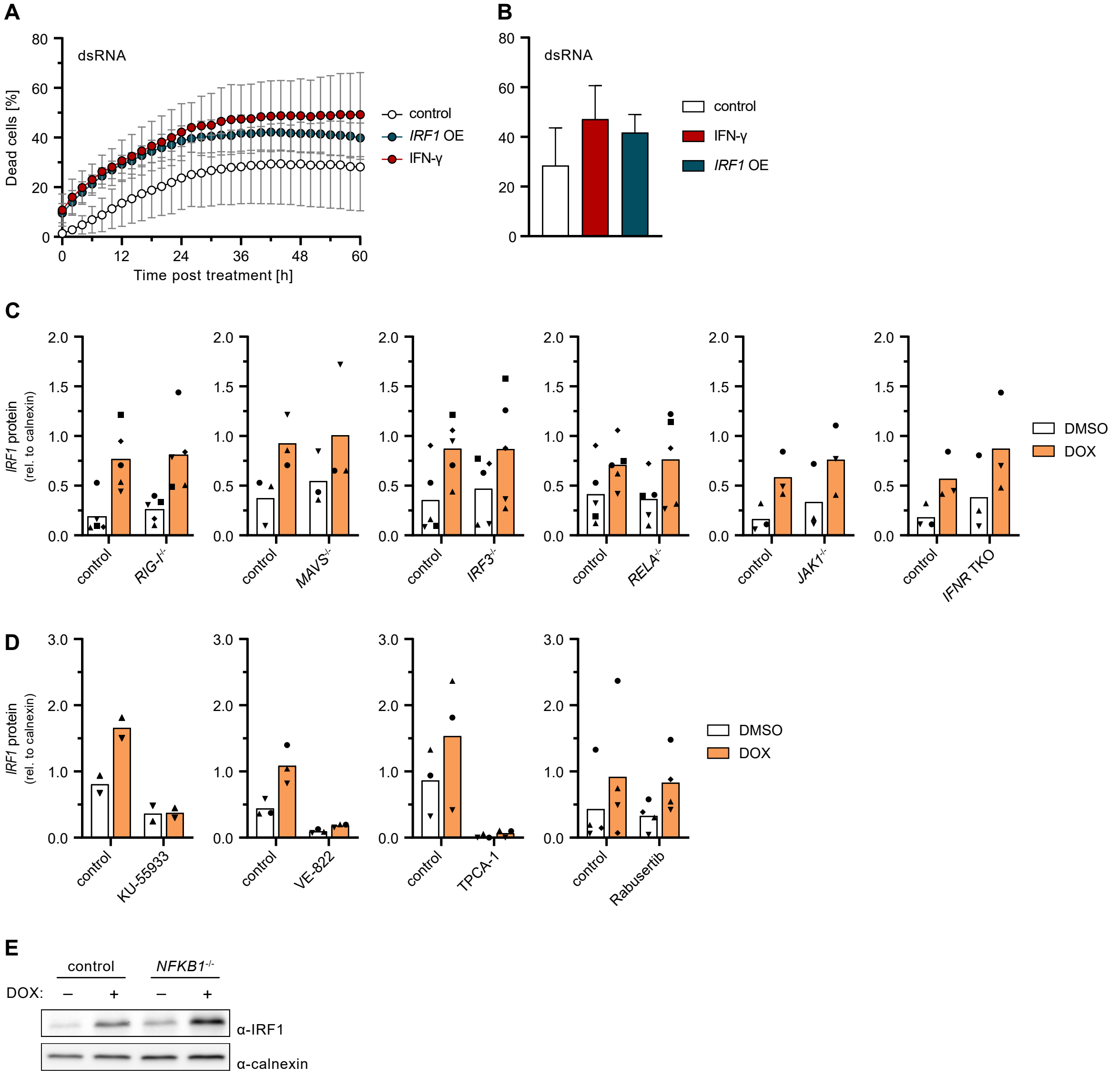


**Fig. S3.**

**(A, B)** Percentage of dead A549 cells with *IRF1* OE or post IFN-γ pre-stimulation relative to total cells counted over time **(A)** or 36 h **(B)** post dsRNA stimulation. **(C-E)** A549 cells with functional KO of the indicated genes or administration of the indicated inhibitors were treated with 2 µM DOX or DMSO for 6 h. Levels of IRF1 were determined by western blot. Signals were quantified and compared to the corresponding calnexin levels (**C, D**). (**A-D**) Data shown represent the results of at least three biologically independent experiments.

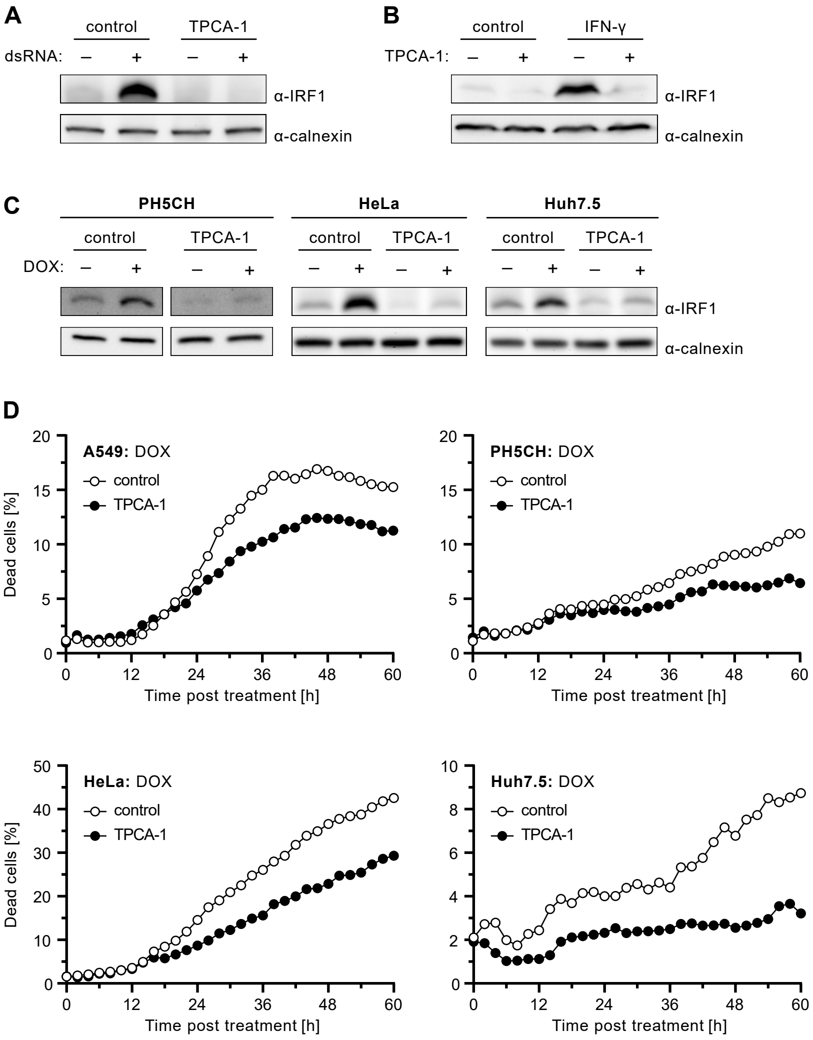


**Fig. S4.**

**(A, B)** A549 cells were treated with TPCA-1, and stimulated with 1.5 ng/ml dsRNA, poly(C), or 200 IU/ml IFN-γ for 6 h. Levels of IRF1 were determined by western blot. **(C)** PH5CH, HeLa, and Huh7.5 cells treated with TPCA-1, and 2 μM DOX or DMSO for 6 h. Levels of IRF1 were determined by western blot. **(D)** Percentage of dead A549, PH5CH, HeLa and Huh7.5 cells relative to total cells counted over time post administration of TPCA-1 and DOX treatment.

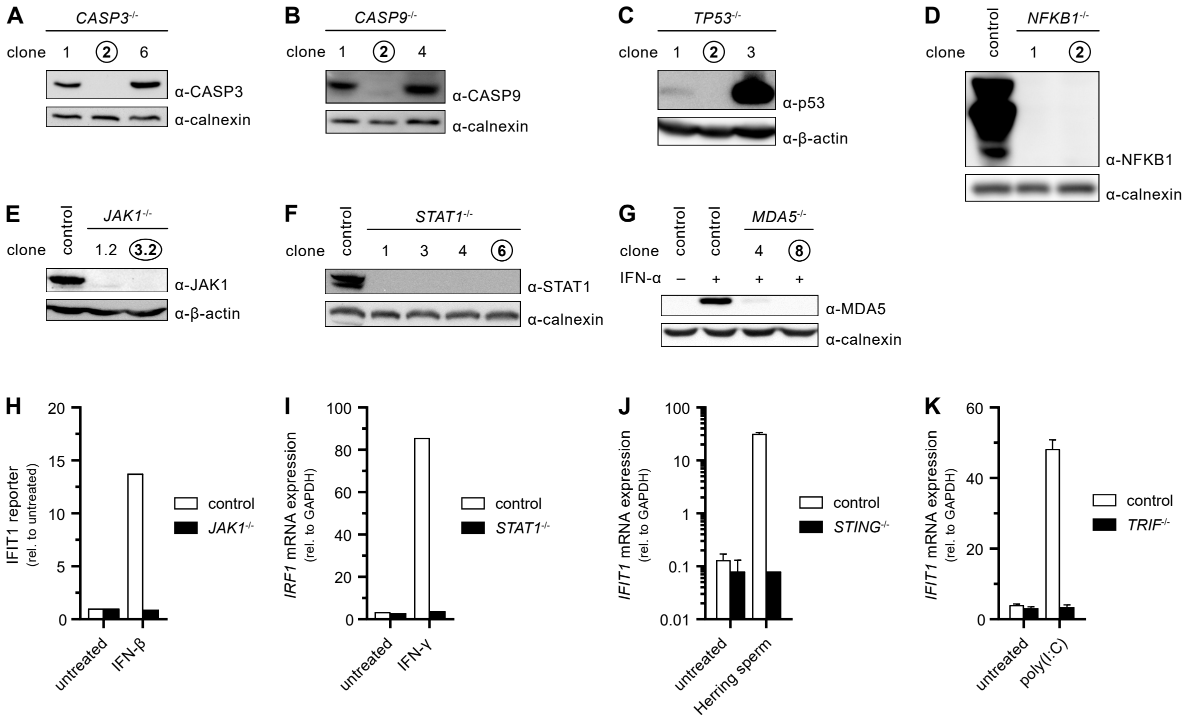


**Fig. S5.**

(**A-G**) Functional KO of the indicated genes in A549 cells was validated by determination of respective proteins by western blot. For *MDA5*, gene expression was increased by pre-stimulation with 200 IU/ml IFN-α (**G**). Circles indicate selected clones with validated KO. (**H**) Gene KO was functionally tested by *Firefly* luciferase-based IFIT1 reporter assay post treatment with IFN-β. Firefly was normalized to Renilla luciferase signal, and values were plotted relative to untreated controls. (**I-K**) Functional KO of the indicated genes was tested by determination of *IRF1* (**I**) *IFIT1* (**J, K**) mRNA transcripts post treatment with IFN- γ (**I**), or Herring sperm DNA and poly(I:C) (**J, K**) by qRT-PCR.
